# Assessing Microbial Diversity in Soil Samples Along the Potomac River: Implications for Environmental Health

**DOI:** 10.1101/2023.03.31.535185

**Authors:** Alexandra Taraboletti, Alexus King, Yasheka Dixon, Oshane Orr, Chevell Parnell, Yasheika Watson, Bruce Nash, Chimdimnma Esimai, George Ude

## Abstract

In this study, we investigated the microbial diversity and community composition of soil samples collected from various sites along the Potomac River within an urbanized region. Our findings revealed the presence of both typical marine soil bacteria and bacterial taxa indicative of urbanization and waste runoff. We observed significant variations in microbial community structure and diversity across different sampling sites, highlighting the influence of environmental factors on microbial abundance and diversity. Through taxon set analysis, we identified shared taxa strongly associated with agricultural pollution, organochlorine pesticide contamination, and bromochloromethane pollution. Additionally, the study revealed potential disparities in human impact, water retention, and tidal/current effects among the soil samples. These insights carry significant implications for understanding the consequences of urbanization on soil microbial communities along the Potomac River and can inform strategies for managing and preserving these ecosystems. Further research is warranted to elucidate the effects of soil health and microbial diversity in this region.

## Introduction

The Potomac River, renowned for its rich historical significance, flows through the Mid-Atlantic region of the United States, traversing the Potomac Highlands before draining into the Chesapeake Bay. The river contains a drainage area of about 14,500 square miles and is approximately 405 miles long. The Potomac basin stretches across parts of four states (Maryland, Pennsylvania, Virginia, and West Virginia) as well as the District of Columbia. The Potomac basin is the 2nd largest watershed in the Chesapeake Bay watershed.^1^ This includes the land areas where water drains towards the mouth of the Potomac, such as the Anacostia River (8.5 miles long), which empties into the Potomac River at Hain’s Point in Washington D.C.^2,3^ The watershed contains a large population, mostly located in the Washington metropolitan area, with forest being the largest land use and agriculture and urban being the second largest land uses in the upper and lower basins respectively.

Human population growth, industrialization, and urbanization have caused a drastic increase in pollution levels within the Potomac basin. ^4–10^ Washington D.C. has a long history of water pollution, with the Potomac and Anacostia rivers subjected to chemical pollution for over two hundred years. The Anacostia River, in particular, is one of the top ten most polluted rivers in the US, containing sewage, metals, PAHs, and PCBs.^9^ Excessive nutrient inputs (mainly nitrogen from nonpoint sources) have caused the eutrophication of surface waters.^11^ The Potomac River is also plagued by high bacterial growth due to sewage runoff and improper waste disposal, leading to the river being used as a sewage drain.^10^ These poor waste management practices and the resulting bacterial growth have led to the formation of cyanobacteria blooms in years of drought and low river volume. The blooms deplete oxygen levels (hypoxic bottom-water dissolved oxygen) and result in the rivers being considered unswimmable and unfishable,^7,10,12^ as confirmed by the Potomac Conservancy (2020), which gave the river a grade of B.^13^ These human-influenced increases in waste and nutrient loads (including discharge from sewage treatment plants, atmospheric deposition, and urban/agricultural runoff) have all negatively impacted the Potomac basin.

In response to these issues, recent initiatives have been launched to maintain the cleanliness and health of the Potomac River, including the Clean Rivers Project, which aims to reduce combined sewer overflows and increase community monitoring of pollutants and toxins.^14^ Sewer separation is just one component of the plan to mitigate combined sewer overflows to the Potomac River and is part of the larger project to clean all three waterways in the District. There has also been a heightened level of community monitoring of pollutants and toxins in the Potomac River, particularly through the efforts of the Interstate Commission on the Potomac River Basin (ICPRB). This increased scrutiny has provided valuable insights into the health of the river and helped to further galvanize support for the protection and preservation of this important waterway.

In line with these efforts, this study aims to investigate the current environmental health of the Potomac River region in the DC metro area via a metagenomic analysis of soil samples from various locations along the basin. Metagenomics, the study of the genetic material of entire microbial communities, has been widely used to investigate the biodiversity of various environments, including soil, water, and air. Soil hosts a wide range of microorganisms that play crucial roles in the ecosystem, and this is especially true for freshwater ecosystems like the Potomac River. The microbial communities present in soil have been shown to be both a marker of and have a significant impact on the overall health of these ecosystems. By examining the metagenomics of soil samples collected along the Potomac River, we can gain valuable insights into the connection between soil microbes and the health of the river basin. The findings of this research have the potential to inform management practices aimed at maintaining and improving the health of the Potomac River for future generations.

## Methods

### Sample Collection

The river soil samples were collected in 50 mL sterile conical tubes, in triplicate, at a distance of 3-5 meters from the banks of the Potomac River and a 6-inch depth from the soil surface. Ethanol and paper towels were used interchangeably between each soil sample collection to ensure clean, sterile tools. Samples were transported to the laboratory within an hour of collection and stored in a -20°C refrigerator until further processed.

### Physiochemical Measurements

Relative nitrogen, phosphorus, and potassium levels were measured using a LaMotte NPK Soil Test Kit. Approximately 0.5 g of soil was extracted for each sample, and the nitrogen, phosphorus, and potassium levels were recorded as specified in the kit procedure. The pH of each sample was collected using a Soil Condition Meter. The soil meter probe was inserted directly into the soil sample and allowed to equilibrate for 1 minute prior to recording the sample pH.

### DNA Extraction

The tubes were ultrasonicated for 1 minute each to achieve cell disruption. Extraction was completed using a Qiagen DNeasy Power Soil kit protocol.^8^ Approximately 0.25g of each soil sample was weighed and recorded. To achieve cell disruption, the samples were ultrasonicated for 1 minute each. Samples were then stored at -20°C until PCR was performed. Nanodrop was used to confirm the quality and concentration of the DNA obtained from the soil samples (Supplementary Table 1).

### PCR/Gel Electrophoresis

DNA products were PCR amplified using primers “515F–806R” targeting the V4 region of the 16S SSU rRNA - used by the Earth Microbiome Project.^9^ Primer sequences are as follows: 515F (Parada)–806R (Apprill), F: GTGYCAGCMGCCGCGGTAA; R: GGACTACNVGGGTWTCTAAT. Primers and primer constructs were designed by Greg Caporaso.^10,11^ Modifications to primer degeneracy were done by the labs of Jed Furhman^12^ and Amy Apprill.^13^ Forward-barcoded constructs were redesigned by Walters.^14^ Amplification conditions were performed in a 25 µl reaction volume and consisted of 13 µL nuclease-free water, 10 µL of 2x PCR master mix, 0.5 µL forward primer, 0.5 µL reverse primer, 1 µL soil DNA. The thermal cycle was programmed for 120 seconds at 94°C as initial denaturation, followed by 35 cycles of 45 sec at 94°C for denaturation, 60 sec at 50 °C as annealing, 90 sec at 72 °C for extension, and a final extension at 72 °C for 10 min. PCR products were examined by gel electrophoresis at 100 V for 45 minutes in a 1% (w/v) agarose gel with ethidium bromide in 1x TAE buffer and compared to a 1 kb DNA ladder (Supplementary Figure 1).

### 16S rRNA Amplicon Sequencing and Analysis

After demultiplexing to assign reads to samples at the New York Genome Center, the resulting FASTQ files were placed in the Cyverse Discovery Environment so that sequences could be analyzed in the DNA Subway Purple Line,^15,16^ which is a graphic user interface for QIIME2. Using the demultiplexed sequence counts summary (Supplementary Figure 2), low-quality sequences were trimmed to position 243 (TruncLenF: 243, TruncLenR: 243). After trimming, samples were rarified (rarification depth Min: 1, Max: 5000), with sampling depths based on the frequency per sample data (Supplementary Figure 3). OTU tables were then generated by matching to the Silva (16S/18S rRNA) database.

### Statistics and Analysis

The Marker Data Profiling (MDP) module in Microbiome Analyst^17,18^ was used to further process the OTU tables generated by DNA Subway. An overview of the library size revealed two sample outliers with <10 read counts (YD1 and CP3; Supplementary Figure 4). These samples were removed from further grouped analyses. To remove low-quality or uninformative features, data filtering was done using a low count filter (minimum count = 4, 20% sample prevalence cutoff) and a low variance filter (10% removed based on inter-quantile range). To deal with the variations in sample depth and the sparsity of the data, normalization (total sum scaling) was applied. Using this data set, Alpha Diversity, Beta Diversity, Interactive Pie, Dendrogram, and Abundance Bar graphs were all generated.

### Data Availability

All metagenomic data is publically available through the Cyverse Platform - DNA Subway: <u>Potomac Soil Metagenomics_2021_2; Project ID: 7799</u>.

## Results

### Sample Collection and Study Site Characteristics

Soil samples were collected in September 2021 from four locations/sites within the Potomac Watershed (Table 1). The four sites, namely, OO, YD, YW, and CP, are each located at different points along the Potomac River, within the boundary of Washington, D.C. (Figure 1). Sampling was performed during a period in which the average temperature was 73°F, and the average rainfall was 0.3 inches (https://www.weather.gov/, Supplementary Figure 5). In this period, the season was characterized as dry; however, for the two days between the collection of the samples, there was light rain in the region.

**Figure 1:**
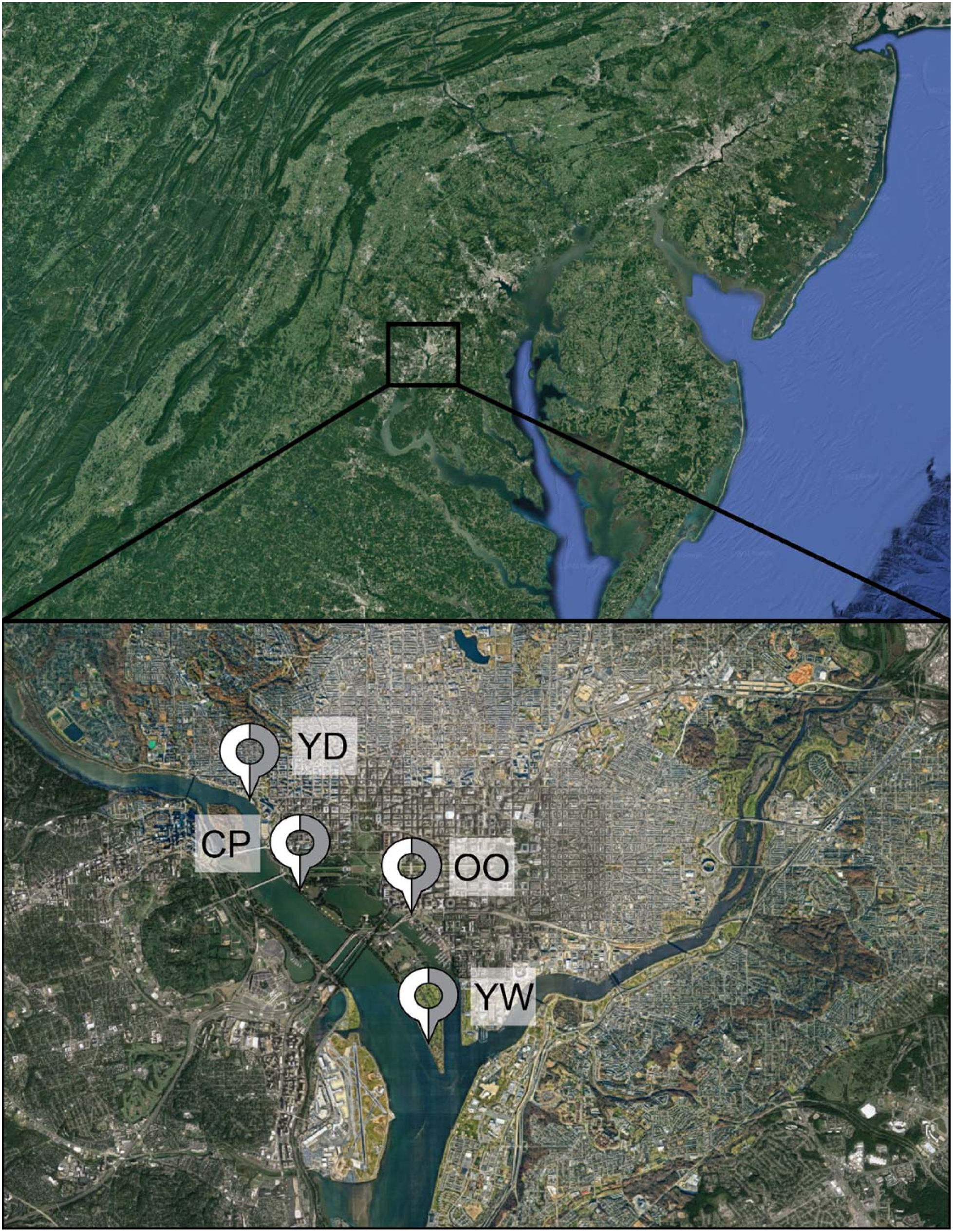
Sample Site Map. Map of Potomac watershed, with DC metro area highlighted. Pins with site names (YD, CP, OO, and YW) show where the Potomac river samples were collected.

**Table 1:** Soil Sampling Sites Along the Potomac River.

| SN# | Date/Time | Site of Collection | GPS Coordinates |
| --- | --- | --- | --- |
| **OO1** | 2:30 PM | SE DC Wharf waterside | Latitude:38.8821 |
|  | 09/22/2021 |  | Longitude: -77.0281 |
| **OO2** | 2:35 PM | SE DC Wharf waterside | Latitude:38.8821 |
|  | 09/22/2021 |  | Longitude: -77.0281 |
| **OO3** | 2:40 PM | SE DC Wharf waterside | Latitude:38.8821 |
|  | 09/22/2021 |  | Longitude: -77.0281 |
| **YD1** | 2:20 PM | Georgetown Waterfront Park | Latitude: 38.900559 |
|  | 09/20/2021 |  | Longitude: -77.058686 |
| **YD2** | 2:26 PM | Georgetown Waterfront Park | Latitude: 38.900559 |
|  | 09/20/2021 |  | Longitude: -77.058686 |
| **YD3** | 2:37 PM | Georgetown Waterfront Park | Latitude: 38.900559 |
|  | 09/20/2021 |  | Longitude: -77.058686 |
| **YW1** | 7:30 PM | 1138-1140 Ohio Dr. SW | Latitude: 38.87792 |
|  | 09/30/2021 |  | Longitude: -77.03797 |
| **YW2** | 7:32 PM | 1138-1140 Ohio Dr. S.W | Latitude: 38.87795 |
|  | 09/30/2021 |  | Longitude: -77.03794 |
| **YW3** | 7:32 PM | 1138-1140 Ohio Dr. SW | Latitude: 38.87795 |
|  | 09/30/2021 |  | Longitude: -77.03794 |
| **CP1** | 2:34 PM | Tidal Basin Washington DC | Latitude: 38.885904 |
|  | 09/22/2021 |  | Longitude: -77.049705 |
| **CP2** | 2:35 PM | Tidal Basin Washington DC | Latitude: 38.885904 |
|  | 09/22/2021 |  | Longitude: -77.049705 |
| **CP3** | 2:36 PM | Tidal Basin Washington DC | Latitude: 38.885904 |
|  | 09/22/2021 |  | Longitude: -77.049705 |

These sites were chosen as they lay along a portion of the river still heavily impacted by combined sewer overflows (CSOs).^19–21^ Washington, DC, has been making progress in cleaning up the Anacostia and Potomac rivers through the court-mandated project known as Clean Rivers.^21^ The project consists of 18 miles of underground tunnels designed to capture CSO before it reaches the rivers. One important tunnel, the “Potomac River Tunnel,” has still yet to be constructed, leaving much of the DC river region – namely the sampled areas - polluted.^19^

### Soil Physiochemical Measurements

The physiochemical measurements (Table 2) of the soil samples collected along the Potomac River revealed that relative nitrogen levels were 20 ppm or below in all samples. The relative phosphorus levels ranged from 4-10 ppm, while all potassium levels were 80 ppm or above. In addition, the pH levels of the soil samples collected along the Potomac River ranged from pH 6.8 to 8.4, with the majority of samples being neutral to slightly basic.

**Table 2:** Soil Physiochemical Measurements.

| SN# | Sample Mass | Nitrogen | Phosphorus | Potassium | pH |
| --- | --- | --- | --- | --- | --- |
| OO1 | 0.501 g | ≤ 20 ppm | ≤ 4 ppm | ≥ 80 ppm | 7.4 |
| OO2 | 0.500 g | ≤ 20 ppm | 6 ppm | ≥ 80 ppm | 7.8 |
| OO3 | 0.501 g | ≤ 20 ppm | ≤ 4 ppm | ≥ 80 ppm | 8.3 |
| YD1 | 0.502 g | ≤ 20 ppm | 10 ppm | ≥ 80 ppm | 8.4 |
| YD2 | 0.500 g | ≤ 20 ppm | 10 ppm | ≥ 80 ppm | 7.7 |
| YD3 | 0.500 g | ≤ 20 ppm | 10 ppm | ≥ 80 ppm | 6.8 |
| YW1 | 0.501 g | ≤ 20 ppm | 6 ppm | ≥ 80 ppm | 7.8 |
| YW2 | 0.500 g | ≤ 20 ppm | 10 ppm | ≥ 80 ppm | 7.3 |
| YW3 | 0.501 g | ≤ 20 ppm | 6 ppm | ≥ 80 ppm | 7.3 |
| CP1 | 0.500 g | ≤ 20 ppm | ≤ 4 ppm | ≥ 80 ppm | 7.7 |
| CP2 | 0.500 g | ≤ 20 ppm | ≤ 4 ppm | ≥ 80 ppm | 8.4 |
| CP3 | 0.501 g | ≤ 20 ppm | ≤ 4 ppm | ≥ 80 ppm | 8.1 |

### DNA Quality and Amplification

DNA samples obtained in the extraction protocol were of good yield (> 20 ng/μL) and quality (260/280 value > 1.8). The measurements of isolated metagenomic DNA for quality and quantity have been presented in Supplementary Table 1. The DNA samples were amplified via PCR targeting the V4 region of the 16S SSU rRNA. Expected amplification was obtained in almost all samples – exhibiting strong bands as visualized with an agarose gel (Supplementary Figure 1). Samples YD3 and CP3 showed weak/no amplification.

### Microbial Diversity and Community Profiling

The amplified metagenomic DNA from these four sites was analyzed via Illumina HiSeq pair-end sequencing, generating 29164 clean reads (average 2430 per sample; range from 1426 to 4367) and 666 operational taxonomy units (OTUs, defined at the 97% sequence similarity). More detailed summary statistics of the sequences can be found in the Supplementary Information. Samples YD3 and CP3, which had reads of 10 and 0, respectively, match samples found not amplified via PCR (Supplementary Figures 1 and 2) and were removed as outliers from further data analysis.

**Figure 2:**
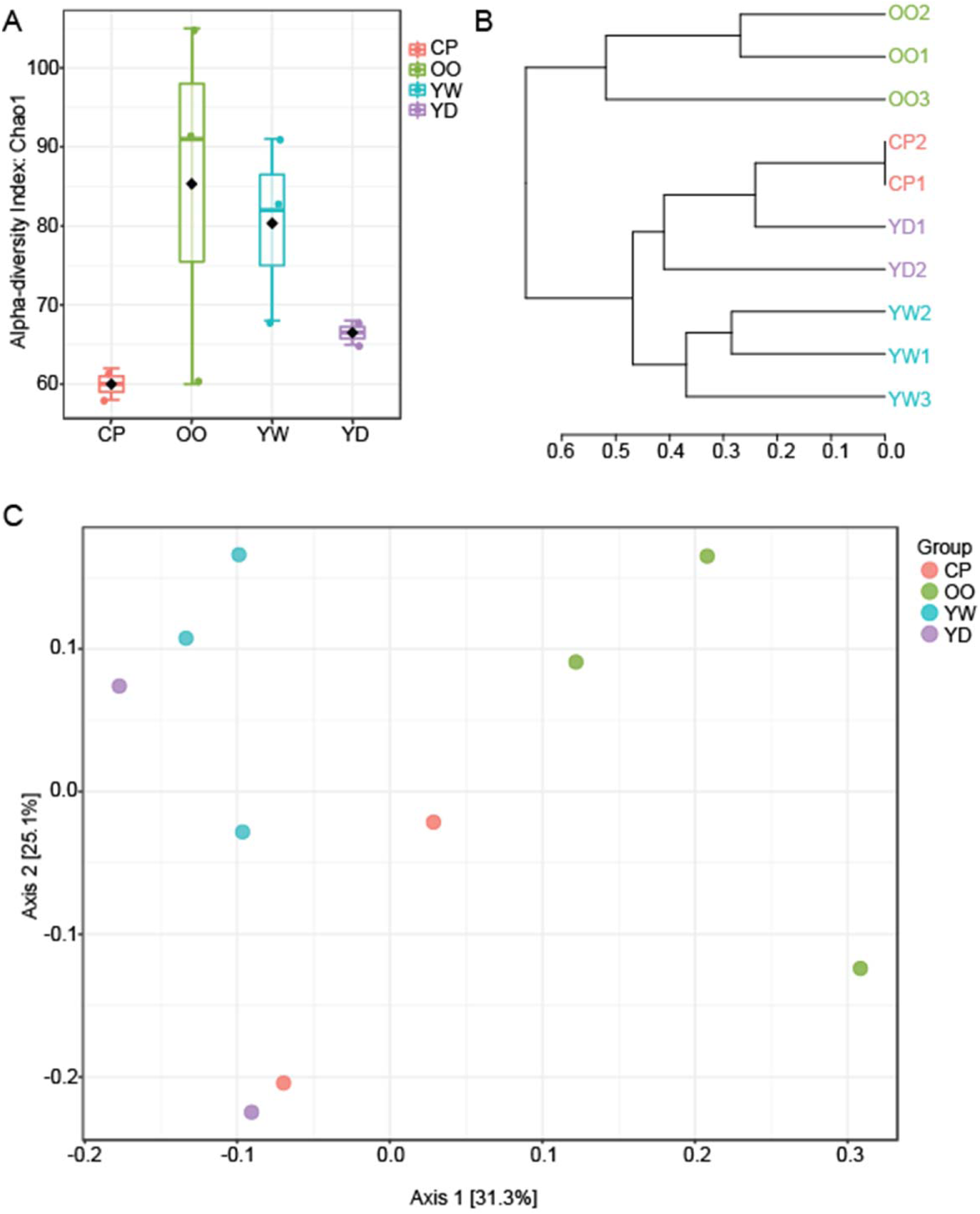
Microbial Diversity. A) Alpha diversity profiling of all samples, utilizing Chao1 diversity measures with ANOVA statistical method. B) Dendrogram analysis of all samples utilizing Bray-Curtis Index distance measures with Ward clustering. C) Beta Diversity profiling of all samples utilizing PCoA 2D ordination plot, with Bray-Curtis Index distance measures and PERMANOVA statistical method. Samples are grouped by sampling site – CP (n=2), OO (n=3), YW (n=3), and YD (n=2).

The distribution of microbial alpha-diversity indices is visualized in Figure 2A. Overall, the mean values in alpha-diversity indices varied among samples grouped by collection site along the river; however, these differences were not statistically significant (Mann-Whitney test, p > 0.05). Notably, communities sampled from a small tidal channel parallel to the main river body (DC Warf; OO) showed higher mean alpha diversity when compared to all other samples. Principal Coordinates Analysis (PCoA, = Multidimensional scaling, MDS) of the normalized OTUs based on Bray-Curtis distances (Figure 2B) also showed a separation between soil samples collected along the main river body (CP, YD, and YW) versus those collected along the parallel channel (OO); this observation is reinforced through Dendrogram analysis (Bray-Curtis distances, Ward clustering; Figure 2C).

Among the identified OTUs, members of bacteria were predominant (98.05%), and a small number of OTUs were classified in the domain of archaea (1.95%). Identification of the OTUs at finer taxonomic levels yielded 73 phyla, 167 classes, 417 orders, 705 families, 1,439 genera, and 570 species. At the phylum level, *Proteobacteria* was dominant (39.0%), followed by *Acidobacteria* (14.0%), *Actinobacteria* (14.0%), *Chloroflexi* (8.0%), *Verrucomicrobia* (7.0%), *Bacteroidetes* (6.0%), *Planctomycetes* (5.0%), *Cyanobacteria* (2.0%), *Gemmatimonadetes* (1.0%), and *Firmicutes* (1.0%); these top 10 bacterial phyla constituted 96% of the total OTUs (Figure 3A). Among *Proteobacteria*, the class *Gammaproteobacteria* was predominant (47.5%), followed by *Alphaproteobacteria* (38.3%) and *Deltaproteobacteria* (14.2%, Figure 3B); among *Acidobacteria*, Subgroup 6 predominates (48.74%), with a noticeable presence of Subgroup 4 (18.75%; Figure 3C).

**Figure 3:**
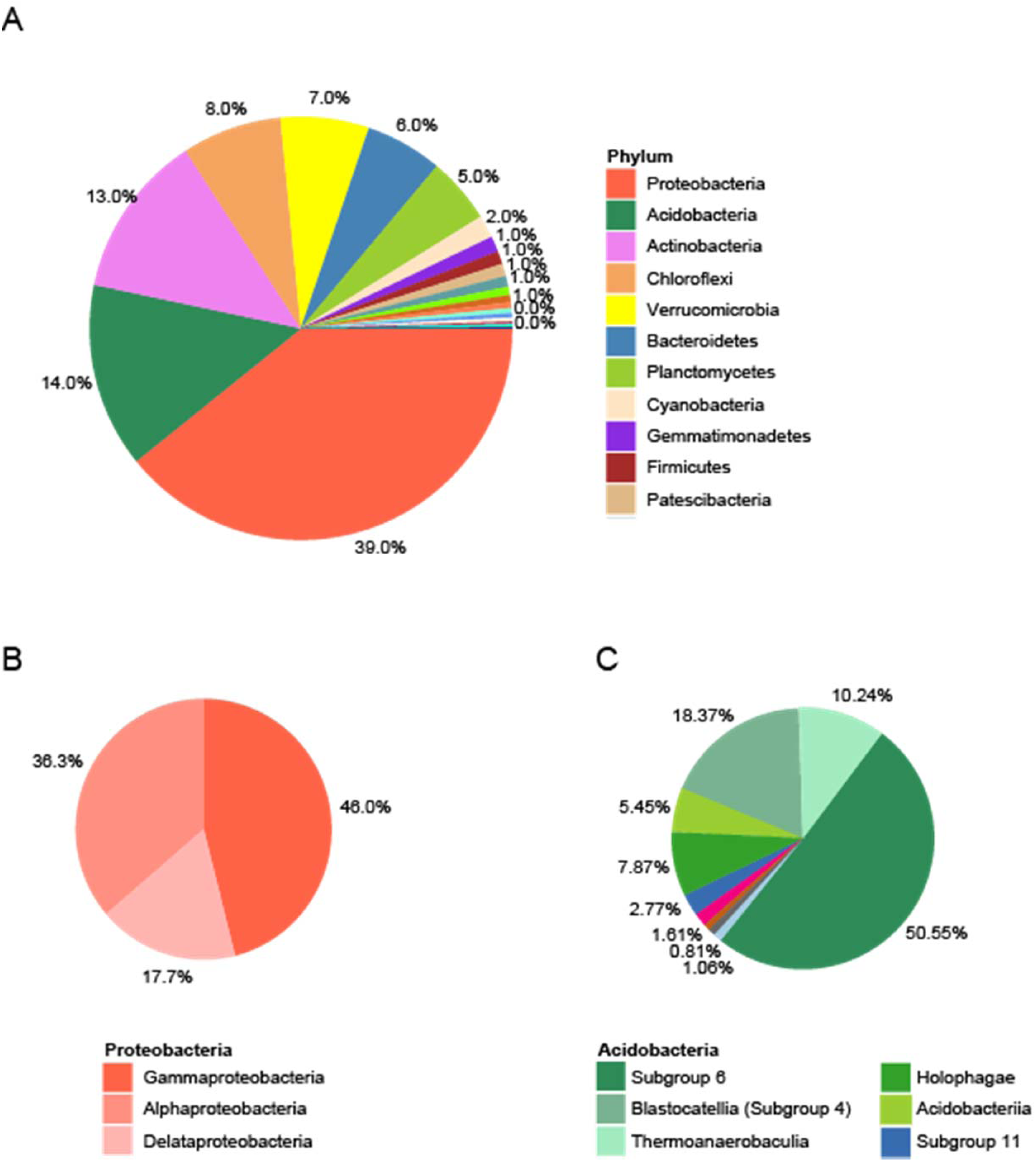
Microbial Community Profiling. A) Pie chart representing the percentage of features found, across all samples (n=10), at the Phylum level. Pie charts of the Class-level composition of the B) Proteobacteria Phylum and the C) Acidobacteria Phylum.

### Microbial Compositions and Group Taxon Analysis

Core microbiome threshold analysis at the family level revealed the top shared/identified taxa to be *Burkholderiaceae, Nitrosomonadaceae, Pedosphaeraceae, Xanthobacteraceae, metagenome, Pirellulaceae, Methyloligellaceae, Pyrinomonadaceae, Gaiellaceae,* and *Solirubacteraceae* (Figure 4A). The Taxon Set Analysis plot (Figure 4B) compares environmental taxon sets to the shared family-level taxa. The plot maps taxa that were highly associated with specific environmental conditions. The shared taxa were found to be significantly associated with agricultural pollution, organochlorine pesticide intoxication, and bromochloromethane pollution. Table statistics can be found in Supplementary Figure 6

**Figure 4:**
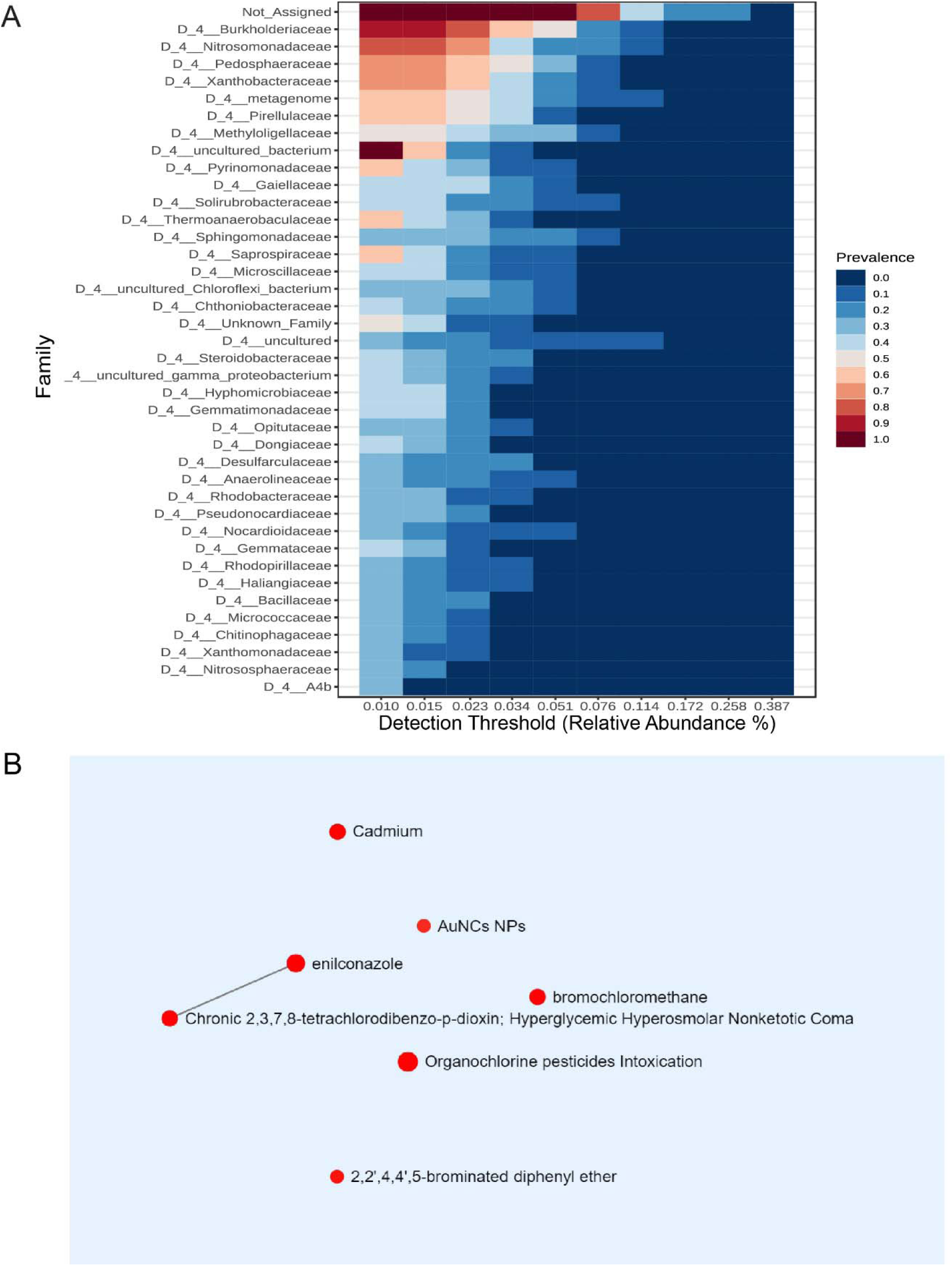
Microbial Compositions and Taxon Set Analysis at the Family level. A) Core microbiome threshold map of the 100 most abundant family level taxa found across all of the sampling sites (n=10). B) Taxon set analysis, enrichment analysis of the top family level taxa found in all samples compared to known environmental taxon set. Each node represents a taxon set, with its color based on its p-value and the size based on the number of matched hits. P<0.05.

### Site-Specific Distribution of Microbial Compositions

Grouped percent abundance data (Figure 5A) highlights taxa abundance differences (genus level) based on the sampling site. Detailed box plot comparisons (Figure 5B) show *Haliscomenobacter, Pseudomonas, Devosia, Luteolibacter, Ilumatobacter, Nitrospira, Steriodobacter,* and *Myxococcales (Blrii41)* were found in higher abundance in location OO compared to the other sites. *Acinetobacter* and *Pseudoxanthomonas* were also found in higher abundance, together at sites OO and CP. *Pirellula, Nocardioides*, *Gaiella,* and *MND1* were found, instead, found upregulated in sites along the upper stretch of the river, sites YW and YD. All correlation tables can be found in Supplementary Table 2.

**Figure 5:**
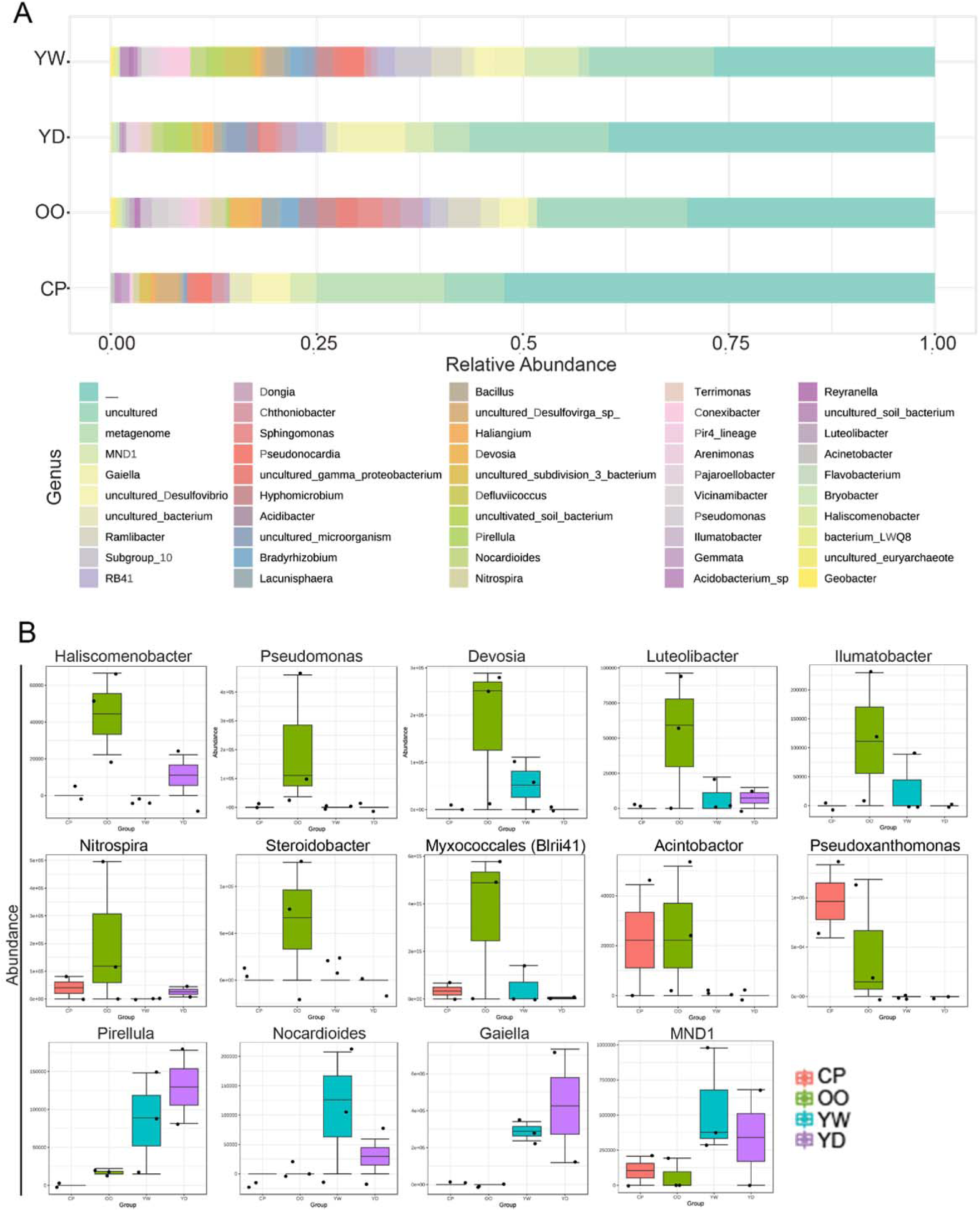
Site-Specific Distribution of Microbial Compositions at the Genus Level. A) Stacked bar plot showing the relative abundance of the top 50 taxa (genus-level) found within sample groups. Samples are pooled by the site collection group. B) Box plots highlighting specific abundance differences in taxa (genus-level). Samples are pooled by the site collection group. CP (n=2), OO (n=3), YW (n=3), and YD (n=2). P<0.05.

## Discussion

The present study reports metagenomic bacteria information, at the phylum and subgroup levels, of soil collected along the DC-metro region of the Potomac River as a means to relay information pertaining to the environmental health of the river basin. Soil collection was conducted at various positions along the river basin (Table 1 and Figure 1), physiochemical aspects of the soil were collected (Table 2), and DNA was extracted and analyzed to gain information about the bacteria communities. 16S bacterial DNA was successfully amplified in all samples (Supplementary Table 1 and Supplementary Figure 1) except for samples YD1 and CP3. This suggests some error in the processing of samples YD1 and CP3, and they should be distinguished as outliers.

Though soil nitrogen, phosphorous, and potassium (NPK) levels can vary depending on the specific environmental context and the types of plants or organisms present – all measured samples had relatively low nitrogen levels, low-medium phosphorous levels, and high potassium levels. Soil samples with low nitrogen levels, ranging from 20 ppm or below, and medium to low levels of phosphorus, ranging from 4-10 ppm, signal limited plant growth and productivity, which could can lead to a decrease in overall biodiversity.^15,16^ The high levels of potassium, above 80 ppm, can indicate that excess fertilizer or manure application has occurred, leading to eutrophication in nearby waterways.^17–19^ The group patterns of the soil microbial data from all samples (Figure 2) displayed variation in alpha-diversity indices among samples but without significant differences (Figure 2A). However, the DC Warf site samples showed higher mean alpha diversity, indicating potential variations in microbial community composition between sites. The PCoA and dendrogram analyses (Figure 2B-C) revealed separation between samples collected along the main river body versus those collected along a small tidal channel, indicating possible discrepancies in human impact, water retention, and tidal/current effects. These patterns also corresponded to differences in soil phosphorous levels, with the samples collected along the tidal channel having slightly lower levels of phosphorous than those collected from the main river (Table 2).

The top taxa shared amongst all sampling sites match typical 16S rRNA-based analyses of phylum diversity found in soil, marine, and wastewater samples (Figure 3).^22,23^ In particular, *Proteobacteria*, *Actinobacteria,* and *Acidobacterium* are well represented and often account for 90% of cultivated soil bacteria (Figure 3A).^22^ *Proteobacteria* is a diverse group of bacteria that is found in a variety of aquatic and terrestrial environments. This phylum is a major contributor to the microbial communities in the Potomac River basin and is likely responsible for the cycling of organic matter, nitrogen, and other essential elements. *Actinobacteria* and *Acidobacterium* are a group of bacteria that are typically associated with soil and are thought to play an important role in the decomposition of organic matter nutrient cycling processes. *Proteobacteria (gamma)*, *Actinobacteria*, and *Acidobacterium (subgroup 6)* (Figure 3B-C) have been reported to thrive in soils with low levels of nitrogen.^20,21^ These findings are consistent with the physicochemical results obtained from the soil samples collected along the Potomac River (Table 2), which showed low levels of nitrogen across all sites. The abundance of these bacterial groups in the soil samples may be attributed to their ability to utilize alternative sources of nitrogen, such as organic matter, or to their capacity to fix atmospheric nitrogen.^20^ The presence of these bacteria is expected, but it confirms that the soil is impacted by human activities.^22,23^ The significant presence of the phylum *Chloroflexi* in the samples may also indicate a shift in the environmental health of the river (Figure 3A). *Chloroflexi* bacteria are often associated with halophilic and thermophilic environments, and their presence suggests that the Potomac river basin could be facing increased hydrological stress. *Chloroflexi* are known to be involved in organohalide respiration and has potential roles in the bioremediation of chlorinated compounds^24^ – noted due to historic PCB pollution in the Chesapeake Bay region.^25–28^ As these bacteria are able to survive in extreme conditions, this could mean that the river basin is subject to increased levels of pollution and other environmental stressors. *Chloroflexi* also plays an important role in activated sludge water treatment plants^29,30,^ and the presence of these bacteria may also be indicative of changes in the river basin’s nutrient cycle. Chloroflexi are known for their ability to break down organic matter, and their presence suggests that there may be an increase in the amount of organic matter entering the river basin.

The identified taxa (family-level; Figure 4A) shared amongst all samples note the strong presence of *Burkholderiaceae, Nitrosomonadaceae*, and *Pedosphaeraceae.* These taxa are indicative of nitrogen cycling and an environment with by-products of sewage and agricultural runoff.^24,25^ *Burkholderiaceae* has been positively correlated with aerobic chemoheterotrophy, aromatic compound degradation, and ureolysis. *Xanthobacteraceae* and *Methyloligellaceae* are typically found in environments with high levels of carbon, which could be due to the abundance of urban settings present along the Potomac River basin.^26–28^ *Pirellulaceae, Pyrinomonadaceae, Sphingomonadaceae,* and *Saprospiraceae* are known to be associated with soil, marine sediments, and biofilms.^29,30^ *Sphingomonadaceae*, *Solibacteraceae*, and *Nitrosomonadaceae* have all been positively correlated with aerobic nitrite oxidation, aerobic ammonia oxidation, and nitrification (all P < 0.05).^31^ The abundance of these bacterial taxa is possibly an indication of elevated levels of urbanization and industrial activity in the vicinity. Furthermore, Taxon Set Analysis (specifically paired to environmental samples) (Figure 4B) suggests that these shared taxa are strongly correlated with agricultural pollution, organochlorine pesticide contamination, and bromochloromethane pollution.

Though many features found in all of the samples were associated with urban settings and pollution, Figure 5 shows that there are differences in taxa abundance found based on sampling sites and that these differences that correspond to sample statistics (Figure 2). For instance, genus-level taxa, such as *Acinetobacter* and *Pseudoxanthomonas,* were more abundant in samples from sites OO and CP, whereas *Pirellula*, *Nocardioides*, *Gaiella,* and *MND1* were more abundant in samples from sites YW and YD. *Haliscomenobacter, Pseudomonas, Devosia, Luteolibacter, Ilumatobacter, Nitrospira, Steriodobacter,* and *Myxococcales (Blrii41)* were overexpressed in samples from site OO. Taxa highly represented from sites YW and YD play a central role in carbon-cycling through methane, and plant and algal degradation, while taxa from sites OO and CP are involved with complex carbon utilization and activated sludge sites. These differences in taxa abundance are likely due to variations in environmental conditions, predominantly the low water current/pooling that occurs at sites OO and CP, as opposed to YW and YD.

## Conclusion

This research provides valuable insights into the microbial diversity and community composition of soil samples collected from various locations along the urbanized stretch of the Potomac River. Our findings underscore the notable variations in microbial community structure and diversity across different sampling sites, emphasizing the influence of environmental factors on microbial abundance. We identified specific bacterial taxa associated with high levels of urbanization, waste sites, and agricultural pollution. Additionally, the study brings attention to potential disparities in human impact and tidal/current effects among the soil samples. These findings carry significant implications for the management and preservation of these ecosystems and contribute to a better understanding of the complex interplay between urbanization and soil microbial communities along the Potomac River. Further research is warranted to more comprehensively explore the impacts of soil health and microbial diversity in this region, with the aim of informing effective strategies for maintaining and improving the health of this vital waterway for future generations.

## Acknowledgments

We gratefully acknowledge the financial support provided by the National Science Foundation (NSF) through Grants 1622811, 183365, and 1438902, which enabled the successful execution of this research project. We would also like to express our sincere gratitude to the Bowie State University Metagenomics CURE Workshop (NSF 1438902), which served as the inspiration and funding source for much of this project. Additionally, we are grateful to the University of the District of Columbia (UDC) STEM Center (NSF 1622811 and 183365) for their unwavering support of our work with undergraduate research students, who played a crucial role in the data collection and analysis for this study. Finally, we thank the various institutions and organizations that facilitated sample collection and provided essential resources, in particular Cold Spring Harbor and the New York Genome Center, ensuring the smooth progress of our investigation.

